# Integration and segregation in the brain as a cognitive flexibility during tasks and rest

**DOI:** 10.1101/2022.10.27.514042

**Authors:** Katerina Capouskova, Gorka Zamora-López, Morten L Kringelbach, Gustavo Deco

**Affiliations:** Center for Brain and Cognition, Computational Neuroscience Group, Universitat Pompeu Fabra, Barcelona, Spain; Department of Psychiatry, University of Oxford, Oxford, United Kingdom; Center for Music in the Brain, Department of Clinical Medicine, Aarhus University, Aarhus, Denmark; Centre for Eudaimonia and Human Flourishing, Linacre College, University of Oxford, Oxford, OX1 2JD, United Kingdom; Department of Neuropsychology, Max Planck Institute for Human Cognitive and Brain Sciences, Leipzig, Germany; Institució Catalana de Recerca i Estudis Avançats (ICREA), Barcelona, Spain; Turner Institute for Brain and Mental Health, Monash University, VIC, Melbourne, Australia

**Keywords:** Integration, Segregation, Brain states, fMRI, HCP data set, Latent Space, Entropy, Classification

## Abstract

To flexibly respond to a continuously changing environment, the human brain must be able to flexibly switch amongst many demanding cognitive tasks. The flexibility inside the brain is enabled by integrating and segregating information in large-scale functional networks over time. In this study, we used graph theory metrics prior to clustering to identify two brain states, segregated and integrated, in 100 healthy adults selected from the Human Connectome Project (HCP) dataset at rest and during six cognitive tasks. Furthermore, we explored two-dimensional (2D) latent space revealed by a deep autoencoder. In the latent space, the integrated state occupied less space compared with the segregated state. After binning the latent space, we obtained entropy from the probability for each data point of being in the bin. The integrated state showed lower entropy than the segregated state, and the rest modality showed higher entropy in both states compared with tasks. We also found that modularity and global efficiency are good measures for distinguishing between tasks and rest in both states. Overall, the study shows that integration and segregation are present in rest and in task modalities, while integration serves as information compression and segregation as information specialisation. These characteristics ensure the necessary cognitive flexibility to learn new tasks with deep proficiency.

## 1. Introduction

When one is confronted with the multimodality of one’s environment, the human brain must flexibly respond and even be prepared when resting [1]. With this constraint, the human brain has evolved into a network with semi-specialised units which are activated when needed and whose activations are integrated in the most connected parts called hubs [2–4]. These structural properties gave rise to cognitive specialisation, when one can properly execute a specific task, and flexibility, when one can easily switch between executing different tasks at once [5–8]. In changing cognitive demands, the human brain must effectively engage in multiple tasks with the same brain structure, which can, however, give rise to different functionalities by adopting varying functional networks [9–11]. When the human brain is using its semi-specialised areas, it segregates received information from the environment; on the other hand, to make sense of the multiple sensory stimuli, it employs integration [12–14]. In this sense, the brain is segregating information to gain specialisation and integrating information to gain flexibility in its cognitive processing [15].

Previous studies [16–18] have proposed that the human brain operates in a mode of metastable equilibrium between local network segregation and global integration of information to flexibly respond to cognitive demands on individuals. In recent neuroimaging studies, modularity and other graph theory metrics have been used to study the organisation of functional networks alternating between integration and segregation in rest condition [19, 20] and during tasks [21–23]. Integration as a flexible adaptation to changing task rules was associated with changes in the global functional connectivity pattern of the fronto-parietal brain network (FPN) [24]. On the contrary, improved segregation from other control networks was found in visual and motor areas due to long-term learning, indicating enhanced independence of basal sensory and motor networks [21].

In this study, we examined integration and segregation in the human brain whilst performing six cognitive tasks and at rest from the HCP dataset. We defined integration as a process when information flows from many nodes to one and is therefore represented as a compression of the signal to gain flexibility. Compressed data need less storage space and less energy to be transferred, so they are easily manipulated, which gives rise to the flexibility in diverse cognitive activities [25]. Segregation, on the other hand, is a projection of information to specialised subnetworks, and here the relationship can be the one-to-many projection of an information flow or a one-to-one projection (Fig. 1H). In our analysis setting, the processes of integration and segregation are presented in a timeframe which we call a state [26–28].

**Figure 1:**
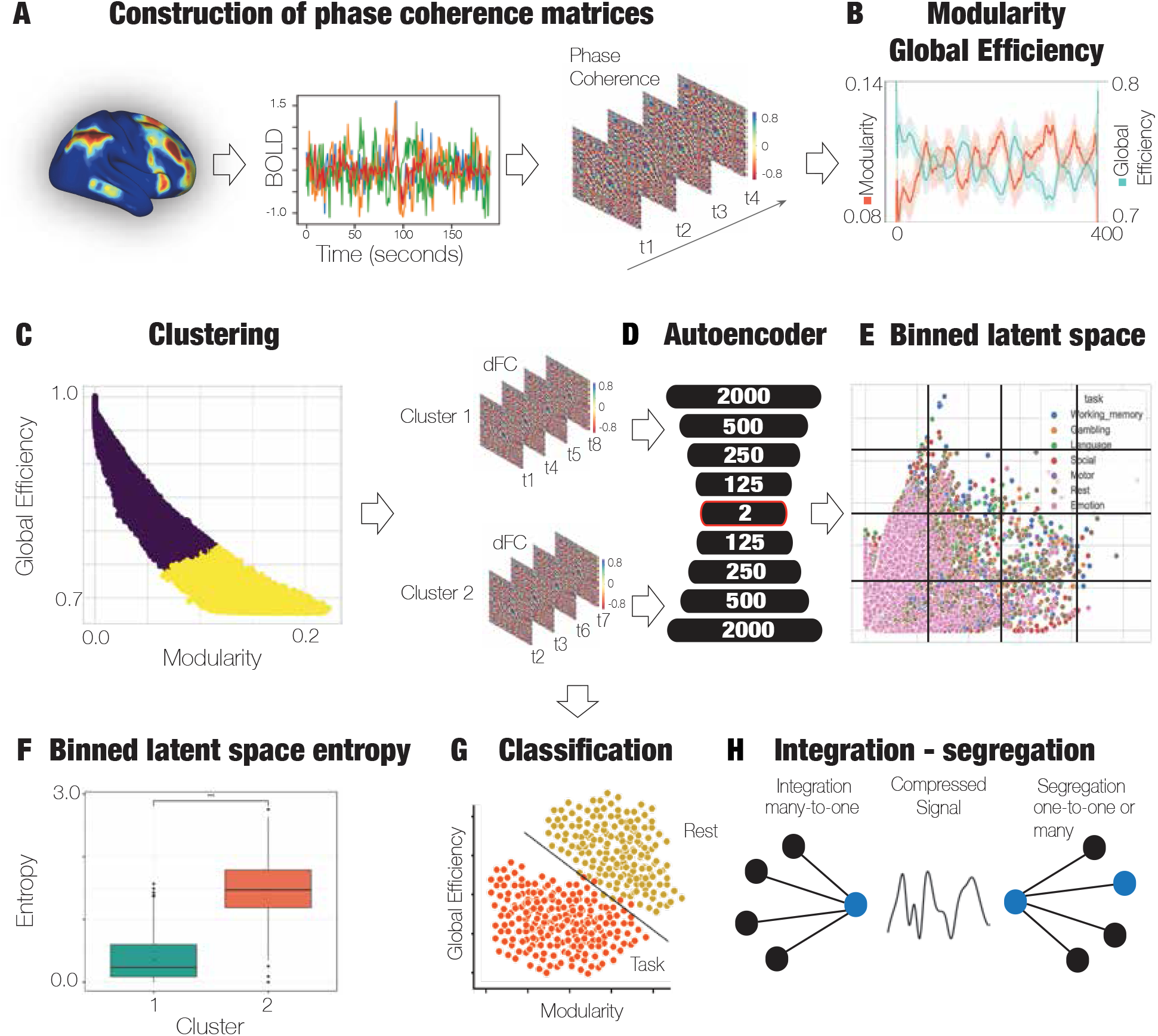
General integration-segregation detection analysis representation (A) BOLD signals from one subject at various brain areas (N = 80). BOLD signals are further filtered with a second-order Butterworth filter and converted into phases of a BOLD signal via the Hilbert transform, and with BOLD phase coherence connectivity, dFC is computed. At each time point t, the cosine function of the BOLD phase difference between brain area b1 and brain area b2 yields asymmetric NxN dFC (b1, b2, t) matrix. (B) Estimation of modularity and global efficiency for each dFC for each subject and modality. (C) K-means clustering model dividing dFCs into two states according to modularity and global efficiency. Division of dFCs is according to the cluster belonging for each time point. (D) Deep autoencoder with 10 layers (nine hidden and one outputlayer). (E) 2D latent space as encoded in the autoencoder and binned into a predefined number of bins. (F) Entropy calculated from probability of being in a certain bin of the latent space. (G) Classification of tasks and rest in the two states as defined by clustering. (H) General schema of integration and segregation.

We used modularity and global efficiency as a measure representatives of integration and segregation in networks inspired by Cohen and D’Esposito [23] (Fig. 1B). We hypothesised that integration is a state with compressed information in the brain, and as such, in latent space with fewer dimensions, it would occupy less space compared with the segregated state. The latent space representation tells us how similar data points are whilst they are compressed. The more similar they are, the less latent space they occupy, and the easier it is to predict where new data points would be. This easier compressibility makes them more predictable with lower entropy. Hence, we also hypothesised that the integration state has lower entropy compared with the segregated state and that in the segregated state, classification of different task modalities and rest would be more accurate compared with the integrated state because of specialisation in segregation.

To assess the hypotheses, we used a computational fMRI data analysis with application of a neural network to achieve lower dimensions for a better inquiry. We applied phase coherence on pre-processed fMRI data which have been converted into phases with Hilbert transform to obtain a dynamic functional connectivity (dFC) matrix (NxNxT, where N stands for the number of brain areas, and T is the number of recorded time points) for each time point (Fig. 1A). To cluster for segregated and integrated states, we used modularity and global efficiency measures, followed by an autoencoder to get a latent space representation of the two states (Fig. 1D). From the 2D latent space, we calculated entropy of the two states (Fig. 1F), and lastly, we included a classification in the states (Fig. 1G).

## 2. Methods

### 2.1. Ethics

All participants gave their full informed consent prior to the study, following the Washington University–University of Minnesota (WU-Minn HCP) Consortium research manual and ethical guidelines. In addition, the study was accredited by the Washington University review board.

### 2.2. Participants

For this study, a representative dataset of 1,003 participants was picked from the March 2017 public data release from the Human Connectome Project (HCP). The subsequent choice of a down-sampled dataset selected 100 unrelated participants (54 females, 46 males, mean age = 29.1+/-3.7 years). The second selection ensures the participants are not related, an important benchmark for omitting potential detected confounds. This also excludes the necessity of accounting for family-structure co-variables in the study.

### 2.3. Neuroimaging acquisition for fMRI HCP

A Siemens 3-T connecome-Skyra scanner was used to scan all the 1,003 HCP participants. In two planned scanning sessions, one session was comprised of working memory, gambling, and motor tasks. The second session included language, social cognition, relational-processing, and emotion-processing tasks. In one session, participants were scanned during rest whilst looking at a projected bright cross on a dark background for approximately 15 minutes. The full dataset with all specifics about the subjects, the scanning protocol, and detailed pre-processing of the data for all cognitive tasks and rest sessions can be viewed at the HCP website (http://www.humanconnectome.org/).

### 2.4. HCP tasks

For the purpose of the HCP study data, seven cognitive tasks were chosen: working memory, motor, gambling, language, social, emotional, and relational, which are explained in greater detail on the HCP website [29]. The tasks were selected to activate particular brain regions associated with distinct cognitive and affective processes. The activations were selected across neural systems like cortical and subcortical and with high sensitivity across subjects. We filtered the six most distinct tasks from the set of HCP tasks. There is a shape identification present in the relational-processing task, which is similarly tested in the emotion-processing task.

### 2.5. Parcellations

For parcellations, commonly used atlases were utilised to prepare all neuroimaging data with the addition of subcortical regions. To achieve a less refined parcellation, the Mindboggle-modified Desikan-Killiany parcellation [30] was adopted (in comparison with the Glasser parcellation [31], because in the study extra dimensionality reductions were processed). The final parcellation comprised of a total of 62 cortical regions, 31 regions per hemisphere [32]. There we also 18 subcortical regions added, nine regions per hemisphere: hippocampus, amygdala, subthalamic nucleus (STN), globus pallidus internal segment (GPi), globus pallidus external segment (GPe), putamen, caudate, nucleus accumbens, and thalamus. With the added subcortical regions, a final parcellation was achieved consisting of 80 regions in the DBS80 parcellation, with a precise demarcation in the common HCP CIFTI ‘grayordinates’ standard space.

### 2.6. Pre-processing and extraction of functional time series in fMRI resting state and task data

A comprehensive explanation of the pre-processing of the HCP resting state and task datasets can be viewed on the HCP website (https://github.com/Washington-University/HCPpipelines/wiki/Installation-and-Usage-Instructions). To concisely outline, the HCP preprocessing pipeline was utilised for both the rest and task data, which apply standardised methods adopting the FMRIB Software Library (FSL), FreeSurfer, and the Connectome Workbench software [33, 34]. The pre-processing pipeline included a correction for spatial and gradient deformities and head motion, intensity normalisation and bias field removal, registration to the T1 weighted structural image, transformation to the 2 mm Montreal Neurological Institute (MNI) space, and, in rest, the use of the FIX artefact removal procedure [34, 35]. In addition, in rest data in the pre-processing procedure, head motion noise was regressed out, and structured artefacts were removed by ICA analysis denoising and the FIX method (independent component analysis followed by FMRIB’s ICA-based X-noisifier [36, 37]). Further details about the ICA-FIX method can be accessed via https://github.com/Washington-University/HCPpipelines/blob/master/ICAFIX/README.md. The refined time series of all ‘grayordinates’ are placed in HCP CIFTI ‘grayordinates’ standard space and are accessible in the surface-based CIFTI file for each participant for the resting state and each task. A custom-built Matlab script was programmed for utilising the ft_read_cifti function from the Fieldtrip toolbox [38] to obtain the average time series of all the ‘grayordinates’ in each brain area of the Glasser and DBS80 parcellations, which are defined in the HCP CIFTI ‘grayordinates’ standard space. The pre-processing protocol included a second-order Butterworth filter in the span of 0.008–0.08 Hz to smooth the BOLD signal from both task and rest data. For all the task and rest data, we first selected only 175 time points (corresponding to the lowest number of time points in the dataset—here, the emotion task) to have balanced data for further analyses.

### 2.7. dFC

To account for the underlying time dynamics, we employed a time-sensitive dFC matrix. For its construction, the BOLD phase coherence connectivity was used [10, 39, 40]. The resulting dFC has NxNxT dimensions, where N is the number of brain areas—80 in the applied DBS80 parcellation—and T is the total number of all time points in the neuroimaging recordings for each task and rest. Firstly, we used a second-order Butterworth filter to smooth the BOLD signal. We estimated the phases of the filtered BOLD signals via the Hilbert transform in all brain areas *n*, getting a phase coherence estimation *θ* (*n, t*). The *dFC* is computed as a cosine similarity amongst phases of brain areas *n* and *p* at time *t: dFC(n, p, t)* as follows:

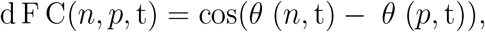

where cos() is the cosine function. The cosine function proved to be beneficial for data scaling as with cos(0) = 1 for areas which occupy a homologous phase for a moment, their BOLD signals are harmonious, and their dFC(n, p, t) is approximately 1. In the case, when two BOLD signals are not consistent with each other, and they are orthogonal, the dFC(n, p, t) is around 0. In the phase coherence estimation, the computation only assigns distances without direction between two phases; hence, the subsequent matrix is undirected. With the characteristic of undirectedness, the matrix is symmetric, and all the relevant information can be obtained from the upper or lower triangular of the matrix.

### 2.8. Modularity and global efficiency

Modularity is a measure informing about the division of network into communities. Before getting the modularity measure (i.e. how well the network is split into groups), it is necessary to run an optimisation algorithm first. In this study, we used the Louvain algorithm for community detection in a weighted undirected graph. Next, we calculated modularity to assess the prevalence of connections inside communities compared with connections between communities. Modularity is defined as follows:

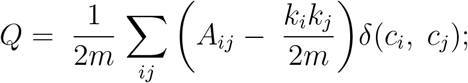

where m stands for the number of edges, A_*ij*_ is the adjacency matrix, k_*i*_ is the degree of i, and *δ* is 1 if i and j belong to the same community and is 0 otherwise.

To obtain a measure of how efficiently a network is trading information between its nodes, we employed a weighted global efficiency measure [41]. Two nodes’ efficiency is the lowest when these nodes are farthest apart. Global efficiency is preferred to path length as is can be used in cases of networks with sparsely connected nodes; parallel processing is also reflected in the efficiency measure. Global efficiency is estimated as the average communication efficiency of a network *G* [42]:

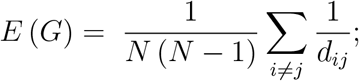

where *d*_*ij*_ is a distance between nodes i and j, or in this case sums of weighted edges traversed, and N is the number of nodes.

### 2.9. Integrated and segregated states—K-means clustering

After estimating modularity and global efficiency for each time point, subject, and condition, we applied an unsupervised machine-learning method to identify the two states: integrated and segregated. There is also an option to select three (or more) states, where one state would serve as a transitional; however, for a good comparison between segregated and integrated, we opted for only two fixed states. We used the k-means clustering method implementation in scikit-learn in Python [43] to cluster for the two states. For quality and validity of the clustering separation evaluation, we applied the silhouette score [44] (Fig 3A). For tolerably separated clusters, the average silhouette coefficient should be at least 0.51 [45].

**Figure 2:**
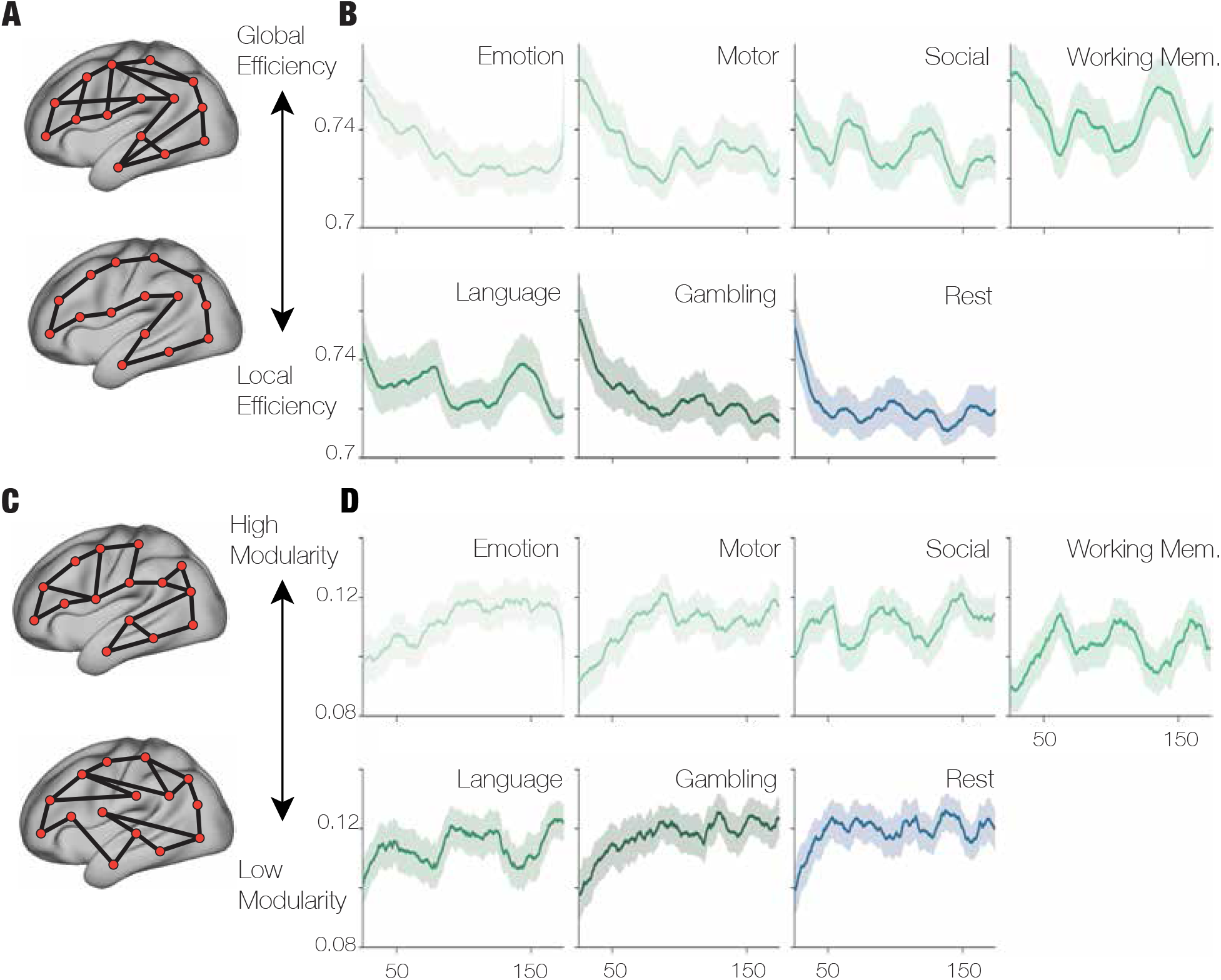
Modularity and global efficiency (A) Illustration of high global efficiency and low global efficiency/local efficiency in a glassmbrain. (B) Global efficiency as detected in all dFCs by task or rest conditions. Oscillating high and low global efficiency was detected in working memory, social, and language tasks associated with high cognitive load of the tasks. In contrast, low global efficiency was detected in emotion tasks, gambling, and rest, indicating lower cognitive loads during these tasks and rest. (C) Illustration of high and low modularity in a glass brain. (D)Modularity as measured in all dFCs by task or rest conditions. Similar results were found with global efficiency, oscillating modularity in working memory, social, and language tasks compared with steady modularity in emotion, motor, gambling, and rest.

**Figure 3:**
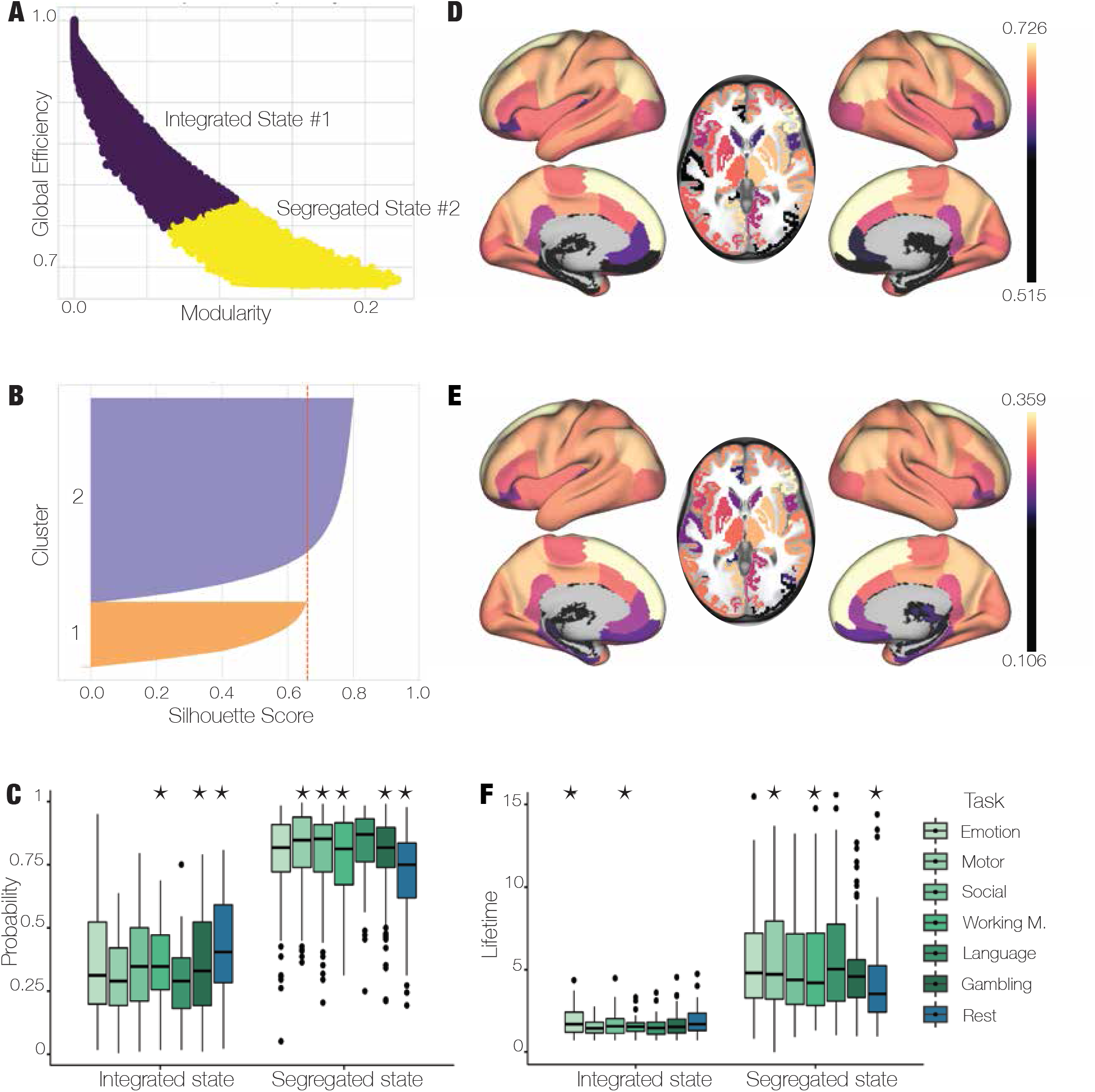
Clustering, brain states, and probability (A) Clustering of modularity and global efficiency into two states, showing that high global efficiency represents the integrated state, and high modularity is associated with the segregated state. (B) Silhouette score of the k-means clustering, which is above the 0.5 threshold and is accordingly well separated. (C) Probability of occurrence in integrated and segregated states: The highest mean probability of being in an integrated state was found in rest, which was also significantly distinct from all the tasks (permutation-based paired t-test, p < 0.01, Bonferroni correction). The highest mean probability of being in a segregated state was found in the language task. Stars indicate a significant difference from all the other conditions (permutation-based paired t-test [p < 0.01] with Bonferroni correction). (D) Brain surface and volume renders of the integrated state. The render represents a normalised connectivity degree in the state. Rendered with Connectome Workbench (available at https://www.humanconnectome.org/software/connectome-workbench). (E) Brain surface and volume renders of the segregated state. The render depicts a normalised connectivity degree in the state. Rendered with Connectome Workbench (available at https://www.humanconnectome.org/software/connectome-workbench). (F) Mean lifetimes of integrated and segregated states in all modalities including rest. In an integrated state, the mean lifetimes are relatively short, especially in motor and language tasks. In a segregated state, the mean lifetimes are longer, with the language task having the longest mean lifetime. Stars indicate a significant difference from all the other conditions (permutation-based paired t-test [p < 0.01] with Bonferroni correction).

### 2.10. Autoencoder

To compare how the integrated and segregated states are represented in space, we had to include a dimensionality reduction method as the dFC matrix of NxN dimensions is too complex to compare with a high number of insignificant features [46]. Although traditional methods such as principal component analysis (PCA) are easy to implement, their linear characteristics would not reveal any interesting structure in the data. For the data representation (the dFCs) in a lower-dimensional space, we implemented a nonlinear data transformation 10-layer-deep autoencoder algorithm. In our case, the latent space is represented only in two features to get a 2D space which is easy to visualise and analyse. We ran one autoencoder for both states at the same time. We also checked if training a balanced dataset (by using oversampling from imbalanced-learn [47]) for both integrated and segregated data produces different results; however, the distinction between the latent space representation in both is still evident. Thus, we opted to use the latent space of an autoencoder trained on unbalanced (real) data points to obtain task and rest differences in the further analyses.

An autoencoder is a special case of an artificial neural network, which is set with random weights trained all at once by minimising the discrepancy between the original data and their reconstruction. An autoencoder is composed of three elements: the encoder, the code, and the decoder. In the first encoder component, the model learns through training to compress the data into an encoded representation. Code is the encoded compressed data representation in a latent space. Lastly, in the decoder, the model is trained to reproduce the input data from the latent space. The reconstruction is optimised to be as close to the original data as possible. The most common training technique is backpropagation, where error derivatives are propagated through the decoder and encoder parts to optimise the weights in the network. The encoder input space (*X*) and decoder input space (*F*) are defined as transitions (*φ, ψ*) which minimize the reconstruction loss so that the divergence between the input data and the output data is reduced and can be defined as follows:

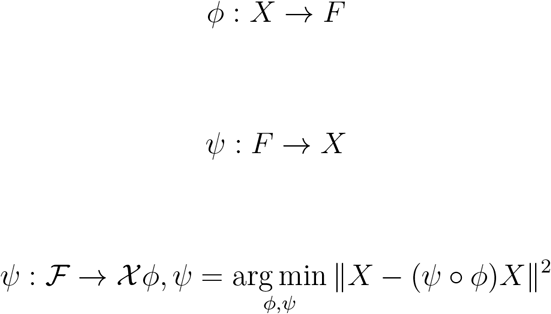

The autoencoder we utilized was structured as a 2000-500-250-125-80-125-250-500-2000 schema inspired by Hinton and Salakhutdinov [48] with the middle layer of latent space that served as the output layer for our further analysis with the encoded vector E(t). We used gradient-based Adam learning rate optimization and applied a mean squared error loss function [49]. In the activation function, the autoencoder ran on rectified linear unit (ReLu) function [50] with 30 epochs (estimated from previous runs as optimal) and batch size of 100.

### 2.11. Binned latent space and entropy

With a latent space of two features, we constructed a 2D space for both integrated and segregated states which we binned (divided space into subsections; Fig. 3.1E) by implementing a function binned_statistics_2D in SciPy [51]. Within the individual bins we counted number of present data points for each condition, subject, and cluster (integrated or segregated). We created a list with each bin’s number of points divided by all the points in the space (also for each subject, condition, and cluster) to count for a data point’s probability of being in the specific bin.

After having a list of probabilities of occurrences in the bins for each condition, subject, and cluster, we calculated the entropy of the occurrence. The entropy is estimated using the following equation:

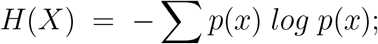

where p(x) is the probability of being in a bin. From the entropy, we obtained an estimate of compression of data in the latent space. The higher the entropy, the less compressed or uniform the occupancy of the latent space.

### 2.12. Classification

We fitted two multiclass support vector machine (SVM) classification algorithms to determine the divisibility of each task and rest based on 80 features (individual measurable properties) taken from the leading eigenvector of each dFC. The two classification models were trained on data clustered as either integrated or segregated. We preferred SVM as it showed a higher performance compared to the k-neighbour and logistic regression in terms of accuracy and overfitting. Furthermore, we utilised a grid search algorithm to search for the optimal parameters and used k-fold cross-validation with five folds to estimate the performance of training and testing on the subset data for the split of 20% test data and 80% training data. We balanced the number of data points for each class according to the lowest occurring number of points: 3282.

### 2.13. Between conditions comparisons

We used a permutation-based paired t-test to discover significant differences between groups. This is a nonparametric two-sample hypothesis test which employs permutations of group labels to estimate the null distribution instead of depending on the test-type standard distributions. The null distribution is calculated independently for each condition. We ran the test for each of 5,000 permutations to compare populations. All reported p-values were corrected with the Bonferroni method for multiple comparisons.

### 2.14. Data and code availability statement

The data for all tasks and rest sessions are publicly available at the HCP website (http://www.humanconnectome.org/). The codes are publicly available at https://github.com/katerinaC/integration_segregation.

## 3. Results

### 3.1. Modularity and global efficiency

First, we estimated graph metrics for all weighted FC matrices for each condition, subject, and time point. We found a handful of statistical differences between tasks and rest in modularity and global efficiency which pass a permutation-based t-test with *p <* 0.01 Bonferroni corrected (Supplementary Table 1). Modularity indicates the structure of a network by its division into communities called modules [52]. High modularity means high connectivity inside communities but low connectedness across the whole network (Fig 2C). This feature indicates higher specialisation with high modularity as we presuppose that each module has its own expertise. In this study, high modularity was achieved in rest, the gambling task, and the emotion task, whilst fluctuating modularity was present in social, working memory, and language tasks (Fig. 2D).

Global efficiency measures how efficiently network’s parts exchange information [42]. With high global efficiency, information is easily transferred in the network, and here we suppose high integration as parts of the brain are easily talking to each other (Fig. 2A). High global functional efficiency was present fluctuating, especially in the working memory task. In social and language tasks, there were also periods with high global efficiency; however, in rest, gambling, motor, and emotion tasks, global efficiency tended to descend during the session (Fig. 2B).

### 3.2. Integrated and segregated states

To get the two states defined as integrated and segregated, we used the k-means clustering algorithm on the two features represented by modularity and global efficiency on all subjects (n = 100) in all tasks and at rest. This separated all dFCs into two groups, and within each group we were able to obtain a mean FC pattern. For the two mean FC patterns, we calculated the normalised degree of connectivity by summation of all the connections in one brain area (in the dFC matrix, a single row) divided by the number of all possible connections for that brain area (number of columns in the dFC matrix). The normalised degree indicates the brain areas dominant in communication which are most functionally connected to the rest areas in the integrated and segregated states.

In the integrated state (Fig. 3D), the most dominantly connected is the left thalamus with the normalised degree of 0.73. Then follow the right superior frontal (0.729), left superior frontal (0.726), right thalamus (0.72), right inferior parietal (0.71), and left inferior parietal (0.71). In the segregated state (Fig. 3.3E), the most connected is the left thalamus with a normalised degree of 0.37; next is the right superior frontal (0.365), followed by the right thalamus (0.36), left superior frontal (0.359), right inferior parietal (0.33), and left inferior parietal (0.33). In general, the most dominant connected brain areas are the same in both states, including the right and left thalami, right and left superior frontal lobules, and right and left inferior parietal lobules.

To examine how often and how long integrated and segregated states are activated, we calculated the average probability of displaying one of the two states in each of the tasks (language, motor, emotion, social, gambling, and working memory) and rest (Fig. 3C). There was a general higher probability of being in a segregated state, particularly in rest; the probability in being in a segregated state was 0.838 ± 0.001 (mean ± standard error of the mean). Rest was followed by the gambling task with a probability of 0.827 ± 0.001, language task 0.805 ± 0.001, and emotion task 0.802 ± 0.001. Slightly less probable was the segregation state in the social task, 0.795 ± 0.001; motor task 0.779 ± 0.001; and least probable in the working memory task 0.72 ± 0.002.

Conversely, the highest probability of expressing an integrated state was in the working memory task 0.453 ± 0.002, emotion task 0.372 ± 0.003, and language task 0.365 ± 0.003. A lower probability of integration was found in the social task 0.364 ± 0.003, motor task 0.356 ± 0.002, and gambling task 0.318 ± 0.003. The least prevalent integration state was revealed in rest, 0.294 ± 0.003.

The longest mean lifetime of a segregated state was measured in the emotion task, 7.255 ± 0.065; followed by the gambling task, 7.155 ± 0.059; and rest, 6.114 ± 0.032. Shorter lifetimes of the segregated state were exhibited in the motor task, 5.726 ± 0.043; emotion task 7.255 ± 0.065; and social task 5.216 ± 0.025. The shortest mean lifetime of the segregated state was identified in the working memory task, 4.401 ± 0.031. In an integrated state, the highest mean lifetime was detected in the emotion task, 2.34 ± 0.044, followed by the working memory task, 1.905 ± 0.01; social task, 1.774 ± 0.013; and language task 1.737 ± 0.013. Relatively shorter mean lifetimes of the integrated state were observed in the motor task, 1.573 ± 0.008; rest, 1.544 ± 0.01; and gambling task 1.542 ± 0.008 (Fig 3F).

The mean silhouette score for the two clusters (0.63) was above the threshold of 0.51 necessary for a reasonably separated clusters within the dedicated space (Fig. 3B).

### 3.3. Autoencoder, binning, and latent space entropy

We trained the autoencoder for 30 epochs, as this number was estimated from previous training sessions as optimal for encoding the data into two dimensions with minimal error in the decoder’s reconstruction (Fig 4A). With 2D latent space, we were able to visualise and analyse the data with the two features, creating a 2D space (Figs. 4B and 4C). Furthermore, we binned the latent space of two features with a different number of bins along each axis. Namely, we used two, four, six, eight, 10, 12, 14, 16, 18, and 20 bins along each axis, creating four, 16, 36, 64, 100, 144, 256, 324, and 400 bins in total. For all those binned latent spaces, we calculated the probability for each subject, condition, and state data point of being in each of the bins. From the probability, we calculated entropy for each of the 10 binned spaces. In each of the 10 binned spaces, entropy of the integrated state was significantly different from entropy of the segregated state with *p <* 0.001 (more in the Supplementary material). We chose the space with 12 bins in each axis to further investigate as it seems to provide balanced space division. This choice provided us with some level of detail in the analysis; however, other choices would also be valid.

**Figure 4:**
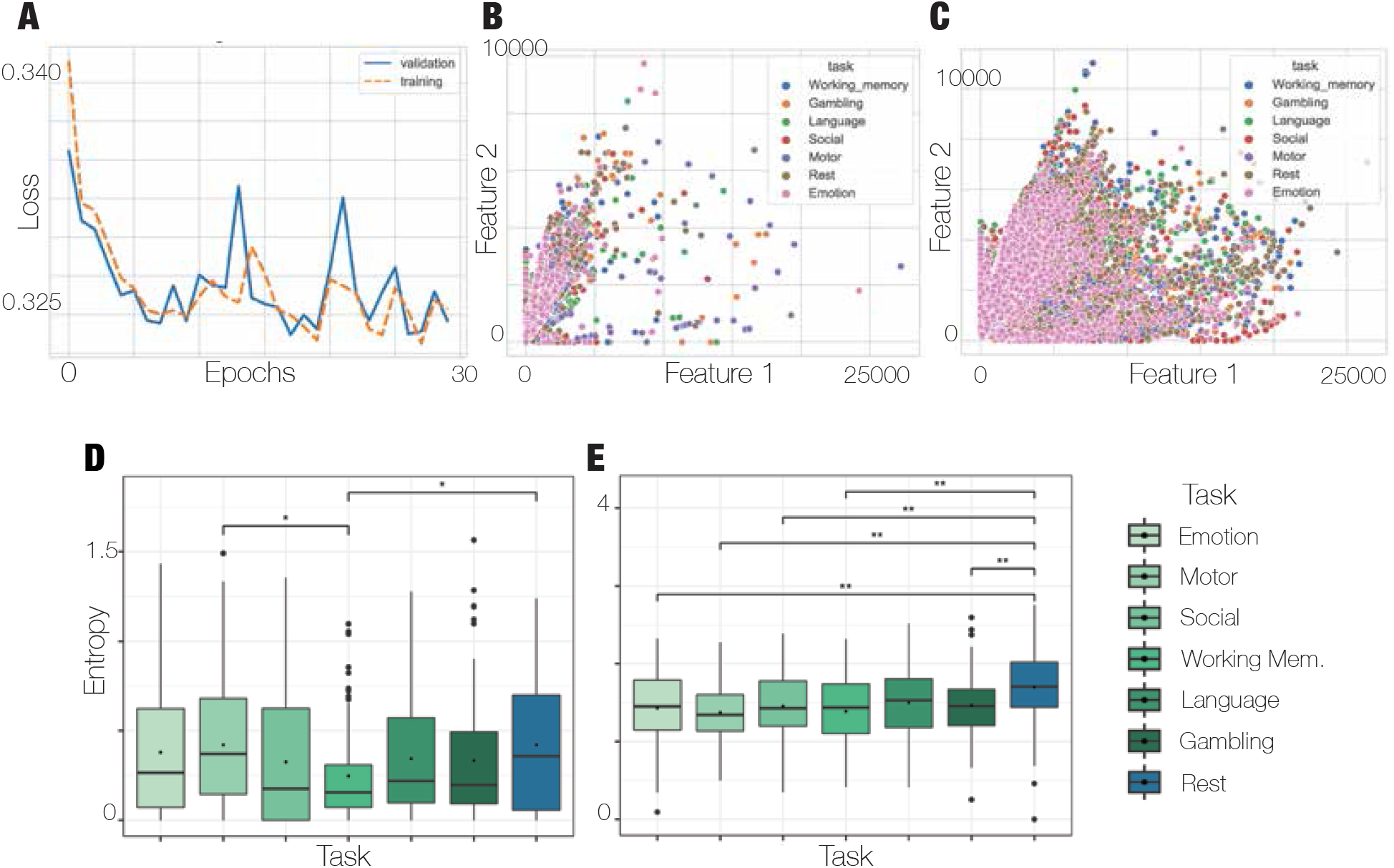
Autoencoder and entropy (A) Loss function for training and validation datasets whilst the autoencoder was trained for 30 epochs. It should be noted that validation is being calculated after the whole batch is trained; hence, that training starts lower. (B) Latent feature space of the integrated state, which is occupied much less compared with the (C) latent feature space of the segregated state, which is very occupied by data points. (D) Entropy of occupation in the integrated state. The highest mean entropy is in the rest condition, with significant differences to the working memory task. Another significant difference is between the motor task and working memory task (permutation-based paired t-test, p < 0.05, Bonferroni correction). (E) Entropy of latent space occupation in the segregated state. Rest shows the highest mean entropy, which differs significantly from all the tasks (permutation-based paired t-test, p < 0.005, Bonferroni correction) except the language task, where p < 0.05 (permutation-based paired t-test, Bonferroni correction).

Integrated and segregated states pass a permutation-based t-test (*t* = − 53.177, *p* = 0.0004) in the 12 bins space scheme. In the same 12-bin separation for both states, we analysed differences amongst all tasks. In the integrated state, only two couples passed a permutation-based t-test with *p <* 0.05, namely motor and working memory tasks (*t* = 4.08, *p* = 0.008) and working memory task and rest (*t* = − 3.88, *p* = 0.008)(Fig. 4D). In the segregated state, there is a higher separability between tasks and rest. Those which passed permutation-based t-tests are the emotion task and rest (*t* = − 4.12, *p* = 0.008), motor task and rest (*t* = −5.4, *p* = 0.008), social task and rest (*t* = − 3.93, *p* = 0.008), working memory and rest (*t* = −53.177, *p* = 0.0004), and gambling task and rest (*t* = −3.77, *p* = 0.02)(Fig. 4E). We also administered separability inside tasks between the two states for the 12 bins space discretisation. For the emotion task, the two states are significantly different (*t* = −18.04, *p* = 0.0004), which is similar to the other conditions: motor task (*t* = −19.11, *p* = 0.0004), social task (*t* = −21.55, *p* = 0.0004), working memory task (*t* = −22.59, *p* = 0.0004), language task (*t* = −19.95, *p* = 0.0004), gambling task (*t* = −21.32, *p* = 0.0004), and rest (*t* = −21.2, *p* = 0.0004) (Fig. 5A).

**Figure 5:**
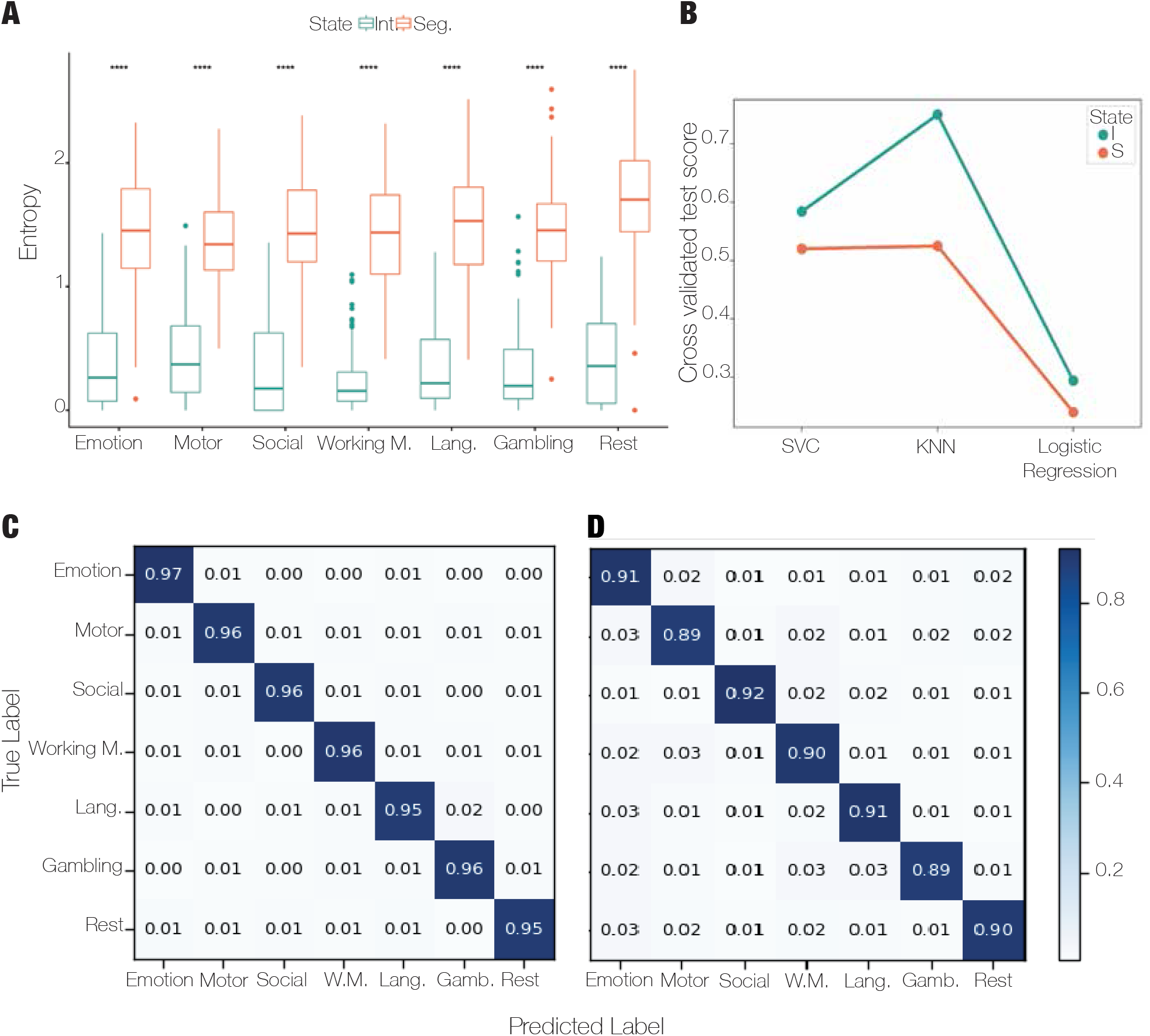
Classification (A) Entropy of latent space occupation in integrated and segregated states for each task. There is a significant difference in all tasks and rest between entropy of latent space occupancy in integrated and segregated states (permutation-based paired t-test, p < 0.0005, Bonferroni correction). (B) Cross-validated test scores for integrated and segregated states in three different classification models:support vector classifier (SVC), k-nearest neighbour (KNN), and logistic regression. The best score was achieved with KNN, which was above 0.5 in both states, and a better score was achieved in the integrated state. (C) Normalised confusion matrix for integrated state prediction in KNN with good prediction results showing on the diagonal, where the label was predicted properly (score above 0.94). (D) Normalised confusion matrix for segregated state prediction in KNN with good prediction results showing on the diagonal, where the label was predicted correctly (score above 0.88).

### 3.4. Classification

After running a several classification methods, we achieved very good separation between rest and the cognitive tasks based on modularity and global efficiency features in the two states. In the integration state, the cross-validated (five-folds) mean test score which the data reached was 0.75 in k-nearest neighbours (compared with support vector classifier [SVC], 0.58, and logistic regression, 0.29; Fig. 5B). The cross-validated mean training score was 0.95 in k-nearest neighbours, 0.77 in SVC, and 0.31 in logistic regression. The most correctly predicted label was in the emotion task (0.97). Other tasks had the same score of 0.95 for correctly predicted labels; rest scored 0.95 (Fig. 5C). In the segregated state, the separation between tasks and rest based on modularity and global efficiency features scored lower than in the integrated state. The cross-validated mean test score in k-nearest neighbours was 0.525 (0.52 in SVC and 0.24 in logistics regression; Fig. 5B). For the training session, the cross-validated mean score was 0.86 in k-nearest neighbours (0.79 in SVC and 0.25 in logistics regression). The most correctly predicted label was in the social task (0.92), followed by emotion and language tasks (0.91). Rest also scored very high in correct label prediction (0.9; Fig. 5D).

## 4. Discussion

This study showed that integration serves as a data compression process to transfer information more efficiently and flexibly. In a 2D latent space which we were able to reach with an autoencoder, integration occupies less space compared to segregation, which was proved by lower entropy in integration compared with segregation. A segregation state is more probable in tasks than in rest as the latter require higher cognitive specificity, and the same is reflected in longer lifetimes of segregated states in tasks.

Inspired by previous studies, we used modularity and global efficiency as indicators of integrated and segregated states [23, 53, 54]; however, other measures could be used from the family of graph theory or those which reflect the characteristics of integration and segregation of information in the brain. In our case, high global efficiency associated with an integrated state means higher communicability amongst distinct brain areas. On the contrary, segregation defined as highly modular demonstrates higher local connectivity and specialisation. We showed that in integration, the enhanced connectivity between distant areas is supported by compressing information and resulting in lower entropy. This is due to the main involvement of structural hubs in the human brain such as the thalamus and superior frontal, which were previously associated with information integration and communication between brain areas [55–60].

We identified high fluctuations between global efficiency / local efficiency and high modularity / low modularity in working memory tasks; this is probably due to the highest cognitive load of the task, indicating the necessity of fast switching and flexibility in cognitively demanding assignments. Similar switching was also demonstrated in social and language tasks. Rest, along with gambling and emotion tasks, generated less switching and a steadier tendency, showing lower cognitive loads.

After training the autoencoder, extraction of its 2D latent space revealed the compactness of occupation of space in the integrated state compared with the segregated state. The integrated state was represented in fewer time points than the segregated state, so we tested whether the compactness could be due to a lower number of data points. We balanced the dataset for training and validating the autoencoder, and the result showed relatively similar results. We also trained the autoencoder more times, and similarly the results showed less space occupation in the integrated state.

After binning the space with different numbers of bins, we chose 12 to analyse further. We calculated entropy of the latent space by estimating the probability for each data point in a particular bin. Highest entropy in both states was in rest. In an integrated state, the difference was significant only from the working memory task; however, in the segregated state, rest entropy was significantly different from all the tasks except language. This shows that the segregated state is driven by specialisation, which is more pronounced in tasks and separates them from rest. Higher entropy in rest was previously identified in other studies [61, 62]. In rest, brain states are characterised by increased metastability, so the brain’s dynamics are less predictable and random whilst visiting a wider dynamical regime. Moreover, in rest, it is harder to predict the state at a particular time point due to higher uncertainty guaranteed by higher entropy [63]. This higher entropy and increased randomness frame allow increased flexibility in learning new tasks with lower precision and specialisation, which is advantageous in settings when no immediate response is necessary—hence, in rest [7, 64, 65]. Rest can be seen as the flexibility generator which provides the edge for the human brain to perform well in specialised tasks but also be able to flexibly obtain new skills.

From our classification results, we can conclude that modularity and global efficiency are good metrics for modalities’ separation. Both states scored high accuracy; however, better results were in the integrated state, which we predicted to have a lower score. This can be due to the data specificity as the segregated state has more data points, which can also be much noisier. The next approach could include three states, where all the middle undecided data points would belong to a special state and not in the segregated one. Some future investigation can also include a dynamical approach [13, 66] to graph analysis and gain more insight into segregation and integration in the brain.

In conclusion, we have identified two states in brain dynamics: integrated and segregated. We have shown that the integrated state occupies less space and is represented with lower entropy compared with the segregated state. With this finding, we proved that integration in this state serves as data compression. We also demonstrated that rest has a higher entropy in both integrated and segregated states compared with tasks and that in the segregated state, the difference is significant between rest and the tasks, pointing towards the specificity in tasks during segregation.

## 5. CRediT authorship contribution statement

Katerina Capouskova: Conceptualisation, Methodology, Software, Validation, Formal Analysis, Investigation, Data Curation, Writing—Original Draft, Visualisation

Gorka Zamora-López: Conceptualisation, Methodology, Writing—Review and Editing, Supervision, Funding Acquisition

Morten L. Kringelbach: Data Curation, Writing—Review and Editing, Supervision, Visualisation

Gustavo Deco: Conceptualisation, Methodology, Formal Analysis, Resources, Writing—Review and Editing, Supervision, Project Administration, Funding Acquisition

## Acknowledgments

K.C, G.Z.-L., and G.D. are supported by the HBP SGA3 Human Brain Project Specific Grant Agreement 3 (grant agreement no. 945539), funded by the EU H2020 FET Flagship programme; G.D. was supported by the Spanish Research Project AWAKENING: using whole-brain models perturbational approaches for predicting external stimulation to force transitions between different brain states, ref. PID2019-105772GB-I00 /AEI/10.13039/501100011033, financed by the Spanish Ministry of Science, Innovation and Universities (MCIU), State Research Agency (AEI). M.L.K. is supported by the ERC Consolidator Grant: CAREGIVING (no. 615539), Center for Music in the Brain, funded by the Danish National Research Foundation (DNRF117), and Centre for Eudaimonia and Human Flourishing funded by the Pettit and Carlsberg Foundations. The funding bodies had neither influence in the research design, data collection, and analysis nor will in preparing and publishing the manuscript.

