## Supplementary materials for "Integration and segregation in the brain as a cognitive flexibility during tasks and rest"

Integration-Segregation Supplementary Material

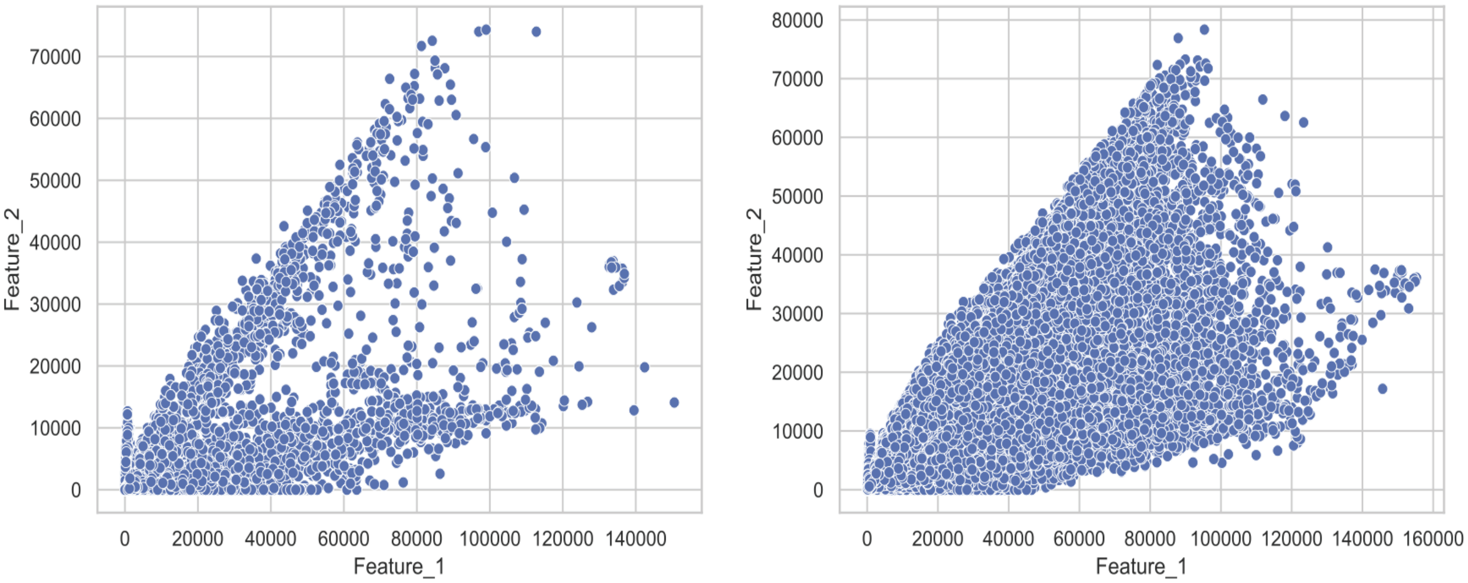


**Fig 1**. 2-dimensional latent space of autoencoder. Latent space of autoendocer that was trained with balanced dataset with the same number of integrated and segregated state data points.

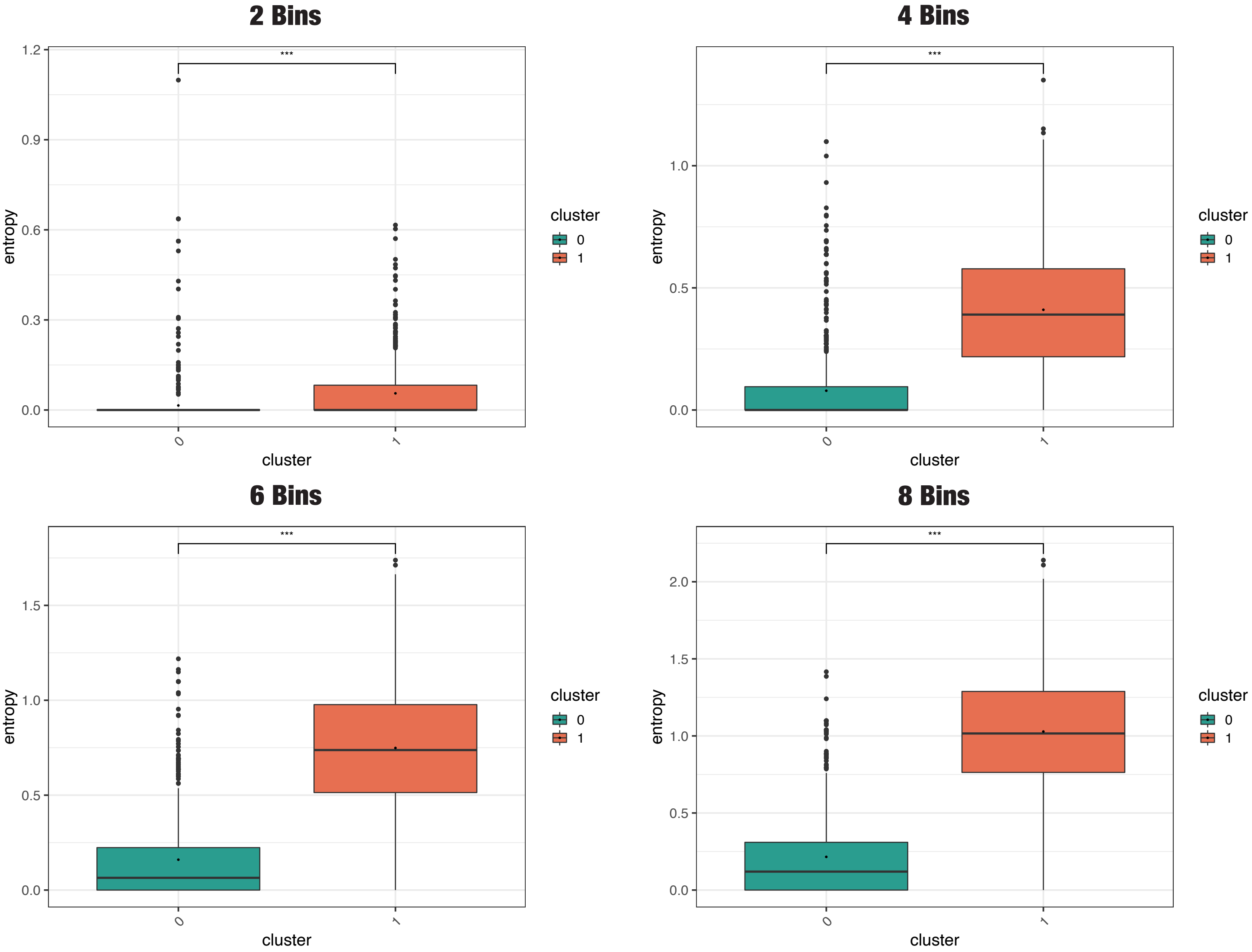

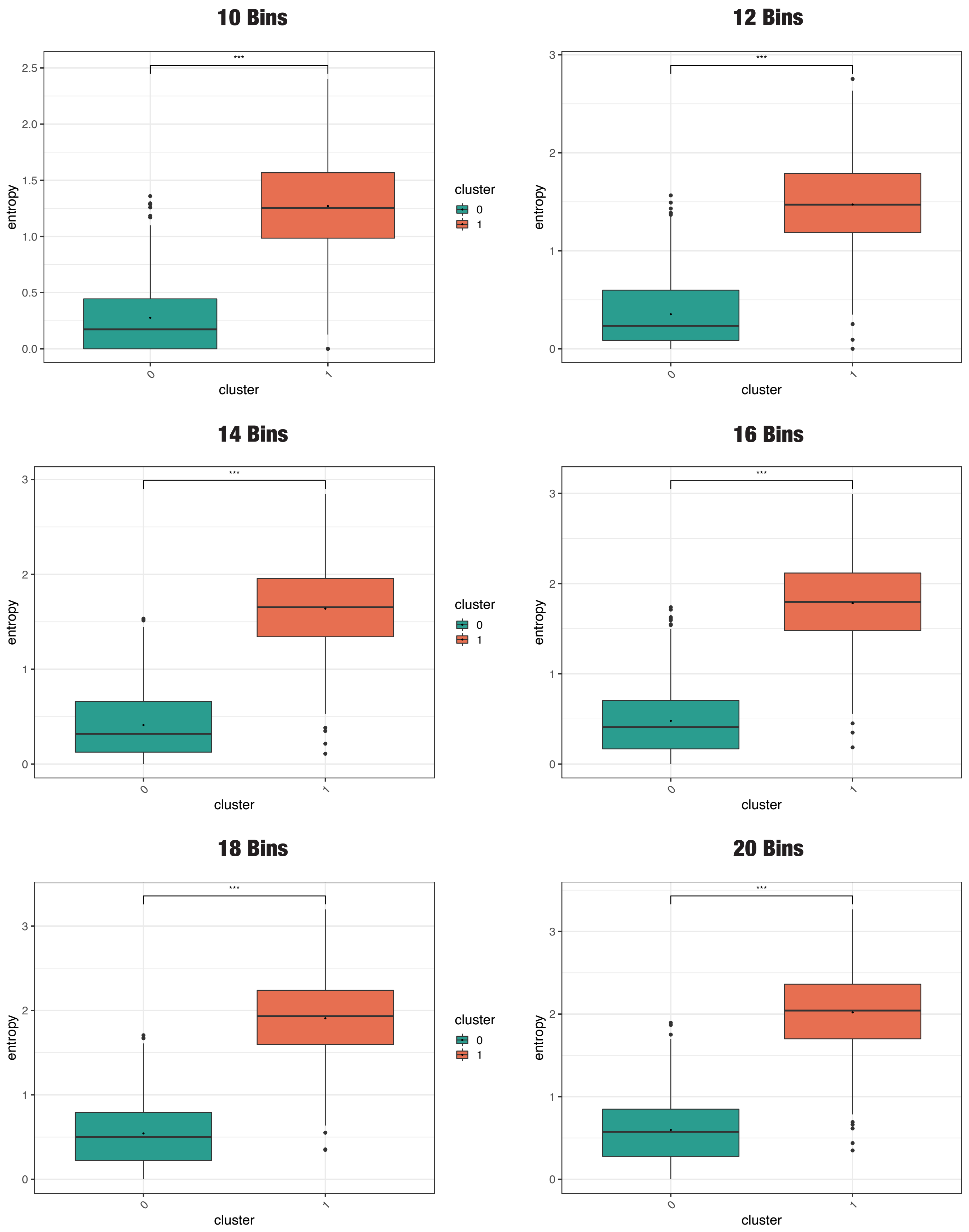


**Fig 2**. Entropy of binned lantent spaces. Latent spaces binned into 2, 4, 6, 8, 10, 12, 14, 16, 18, and 20 bins all show significant differences in entropy between integrated and segregated states (permutation-based paired t-test, $p < 0.005$, Bonferroni correction).

| | Conditions | Modularity p value Bonferroni | Global efficiency p value Bonferroni | Modularity t value | Global efficiency t value |
|---|---|---|---|---|---|
| 0 | Emotion Motor | 0.016796640671865627 | 1.0 | 3.269561914340805 | -0.4352789644699983 |
| 1 | Emotion Social | 1.0 | 0.7810437912417517 | -0.2137229613652199 | 2.083273911581249 |
| 2 | Emotion Working_memory | 0.008398320335932814 | 0.008398320335932814 | 8.63781598135505 | -4.638396212115413 |
| 3 | Emotion Language | 0.008398320335932814 | 0.008398320335932814 | -9.954013862790669 | 11.643818616105715 |
| 4 | Emotion Gambling | 0.008398320335932814 | 0.008398320335932814 | -13.777255800295222 | 13.269594635080477 |
| 5 | Emotion Rest | 0.008398320335932814 | 0.008398320335932814 | -18.579518612049704 | 17.516469839216803 |
| 6 | Motor Social | 0.008398320335932814 | 0.050389922015596875 | -4.006112048426844 | 3.00243787916779 |
| 7 | Motor Working_memory | 0.008398320335932814 | 0.008398320335932814 | 6.150327298550887 | -5.072036994591516 |
| 8 | Motor Language | 0.008398320335932814 | 0.008398320335932814 | -15.581299494817443 | 14.506982210722013 |
| 9 | Motor Gambling | 0.008398320335932814 | 0.008398320335932814 | -19.68821822733607 | 16.32544431161712 |
| 10 | Motor Rest | 0.008398320335932814 | 0.008398320335932814 | -27.738170396379108 | 22.427878498989884 |
| 11 | Social Working_memory | 0.008398320335932814 | 0.008398320335932814 | 10.360571413527477 | -8.140396629919403 |
| 12 | Social Language | 0.008398320335932814 | 0.008398320335932814 | -11.240247317562249 | 11.05799643535622 |
| 13 | Social Gambling | 0.008398320335932814 | 0.008398320335932814 | -15.508577319238311 | 13.009044253487845 |
| 14 | Social Rest | 0.008398320335932814 | 0.008398320335932814 | -22.181906857927792 | 18.150597140774895 |
| 15 | Working_memory Language | 0.008398320335932814 | 0.008398320335932814 | -23.182006248142162 | 20.699224903213835 |
| 16 | Working_memory Gambling | 0.008398320335932814 | 0.008398320335932814 | -26.995416296871365 | 22.08693958216318 |
| 17 | Working_memory Rest | 0.008398320335932814 | 0.008398320335932814 | -40.06722703450974 | 32.4018731795846 |
| 18 | Language Gambling | 0.008398320335932814 | 0.15116976604679064 | -5.0519475155451365 | 2.7689429616244574 |
| 19 | Language Rest | 0.008398320335932814 | 0.008398320335932814 | -8.735534216478994 | 4.808243989623918 |
| 20 | Gambling Rest | 1.0 | 1.0 | -1.8059545343238956 | 1.0312906103264756 |

**Table 1**. P-values from a permutation-based t-test with Bonferroni correction between different tasks and rest in modularity and global efficiency.
